# Identification of ubiquitin variants that inhibit the E2 ubiquitin conjugating enzyme, Ube2k

**DOI:** 10.1101/2021.05.28.446107

**Authors:** A.J. Middleton, J. Teyra, J. Zhu, S.S. Sidhu, C.L. Day

**Author notes:** Author for correspondence Department of Biochemistry, School of Biomedical Sciences, University of Otago, Dunedin 9054, New Zealand.

## Abstract

Transfer of ubiquitin to substrate proteins regulates most processes in eukaryotic cells. E2 enzymes are a central component of the ubiquitin machinery, and generally determine the type of ubiquitin signal generated and thus the ultimate fate of substrate proteins. The E2, Ube2k, specifically builds degradative ubiquitin chains on diverse substrates. Here we have identified protein-based reagents, called ubiquitin variants (UbVs), that bind tightly and specifically to Ube2k. Crystal structures reveal that the UbVs bind to the E2 enzyme at a hydrophobic cleft that is distinct from the active site and previously identified ubiquitin binding sites. We demonstrate that the UbVs are potent inhibitors of Ube2k and block both ubiquitin charging of the E2 enzyme, and E3-catalysed ubiquitin transfer. The binding site of the UbVs suggests they directly clash with the ubiquitin activating enzyme, while potentially disrupting interactions with E3 ligases via allosteric effects. Our data reveal the first protein-based inhibitors of Ube2k and unveil a hydrophobic groove that could be an effective target for inhibiting Ube2k and other E2 enzymes.

## Introduction

Ubiquitin transfer is a post-translational modification that plays a critical role in almost all aspects of eukaryotic cells, including control of protein degradation by the proteasome (Hochstrasser, 1996), modulation of gene expression (Mark and Rape, 2021), recruitment of proteins to signalling platforms (Chen, 2005; Stewart et al., 2016), and dictation of the precise timing of cell division (Pines, 2011). Consequently, dysregulation of ubiquitin transfer results in many diseases, such as cancers, immune disorders, and neurodegenerative diseases (Rape, 2018). As a result, the ability to manipulate the components of the ubiquitin system is of considerable interest for treatment of disease.

Ubiquitin transfer is governed by three families of enzymes: the E1 ubiquitin activating enzymes, the E2 ubiquitin conjugating enzymes, and the E3 ubiquitin ligase enzymes (Gundogdu and Walden, 2019). Together, the machinery covalently links ubiquitin to a substrate Lysine or N-terminal Methionine residue with an isopeptide bond. Ubiquitin itself can be a substrate as it contains seven Lys residues, and this results in the formation of ubiquitin chains with distinct consequences (Osborne et al., 2021). For example, chains linked by Lys63 can act as scaffolds for recruiting proteins to signalling cascades, whereas Lys48-linked ubiquitin chains typically result in degradation of the attached substrate by the proteasome (Osborne et al., 2021). The nature of the ubiquitin signal is typically dictated by the E2 enzyme, of which there are ∼40 in humans. Depending on their structure and biological context, E2 enzymes can add a single ubiquitin moiety or ubiquitin chains of various types to substrate proteins. Because the downstream effects of ubiquitin transfer are entirely dependent on the nature of the ubiquitin signal, E2 enzymes have a central role in ensuring substrate proteins are correctly modified by the ubiquitin machinery.

E2 enzymes are characterised by a conserved ubiquitin conjugating domain (UBC) that interacts with an E1 enzyme and E3 ligase via conserved and partly overlapping interfaces. After charging of the E2 enzyme with ubiquitin, the resulting E2∼Ub conjugate disengages the E1 and interacts with one of the hundreds of E3 ligases. The E3 ligase typically coordinates the choice of substrate to be modified, while also activating the E2∼Ub bond so that it is susceptible to nucleophilic attack (Dou et al., 2012; Koliopoulos et al., 2016; Plechanovova et al., 2012; Sanchez et al., 2016; Scott et al., 2014). When activated, the conjugated donor ubiquitin makes extensive contacts with the E2 enzyme, including nestling of ubiquitin’s flexible C-terminal tail into a shallow groove of the E2 enzyme. When building ubiquitin chains, the E2 enzyme must also interact with a substrate ubiquitin, and this is referred to as the acceptor ubiquitin. Disrupting any of these protein-protein interactions can disable the E2 enzyme.

The ubiquitin conjugating enzyme, Ube2k, produces Lys48-linked ubiquitin chains exclusively, and promotes the degradation of its targets in cells (Chen and Pickart, 1990). A recent report indicates that Ube2k is involved in promoting the degradation of the pro-survival Bcl-2 protein family member, Mcl-1, suggesting Ube2k may have a pro-apoptotic role (Djajawi et al., 2020). Other studies suggest that Ube2k is involved in overcoming cell-cycle arrest in response to DNA damage by promoting the degradation of p53 (Hong et al., 2019). In addition, Ube2k levels contribute to multiple neurodegenerative diseases that are characterised by aggregation of proteins. For example, Ube2k is involved in Huntingtin disease (Kalchman et al., 1996); is associated with cell death from polyglutamine diseases (de Pril et al., 2007); is proapoptotic in response to the accumulation of amyloid-β (Song et al., 2003); and is elevated in the brains of individuals with schizophrenia (Meiklejohn et al., 2019). Recent work suggests that Ube2k deficiency results in motor impairment reminiscent of Parkinson’s disease, and could act as a biomarker of the disease (Su et al., 2018). There is also evidence that Ube2k can add Lys48-linked chains onto already-established Lys63 ubiquitin chains, and thereby quench Lys63-induced signalling (Pluska et al., 2021). While the roles of Ube2k are diverse, they are unified by the ability of Ube2k to promote the degradation of substrate proteins. Development of inhibitors that specifically target Ube2k would not only provide tools to allow a greater understanding of its function, but may also provide a framework for small-molecule design.

Here we have used a phage-displayed ubiquitin variant (UbV) library to isolate specific inhibitors of Ube2k. Of the six UbV binders identified, two are potent inhibitors of ubiquitin transfer promoted by Ube2k. Biochemical experiments show that both UbVs inhibit charging of Ube2k by the E1 enzyme, and also E3-catalysed discharge. Structures of the UbV-Ube2k complexes show that both UbVs bind in a hydrophobic cleft distant from the active site, and this site includes E2-E1 contacts. While this explains why charging of Ube2k is impeded, the UbV binding site does not overlap with the E3-binding site, and the reason for decreased E3-catalysed discharge is less certain. Our research reveals a hydrophobic groove on Ube2k that can bind ligands to block the assembly of degradative ubiquitin chains.

## Results

### Selection and characterisation of UbVs that bind Ube2k

To discover specific modulators of the E2 enzyme Ube2k, we generated a stable Ube2k∼Ub conjugate and performed selection using a highly diverse phage-displayed library containing 2 × 10^9^ unique UbVs. The library was a further iteration of libraries generated for selection against deubiquitinases, E3 ligases, and other E2 enzymes (Ernst et al., 2013; Gabrielsen et al., 2017; Teyra et al., 2019). In this library, residues were diversified across the surface of ubiquitin that is involved in the vast majority of ubiquitin-protein interactions, including contacts with E2s, ubiquitin-associating domains, and deubiquitinases. The surface comprises the beta sheet of ubiquitin and its five flexible C-terminal residues (Figure 1A). To minimise disruptions to the fold of ubiquitin while maximising diversity across the surface, the library was built using a ‘soft-randomisation’ approach with degenerate codons that encode approximately 50% wild-type sequence at each diversified position (Sidhu et al., 2000).

**Figure 1.**
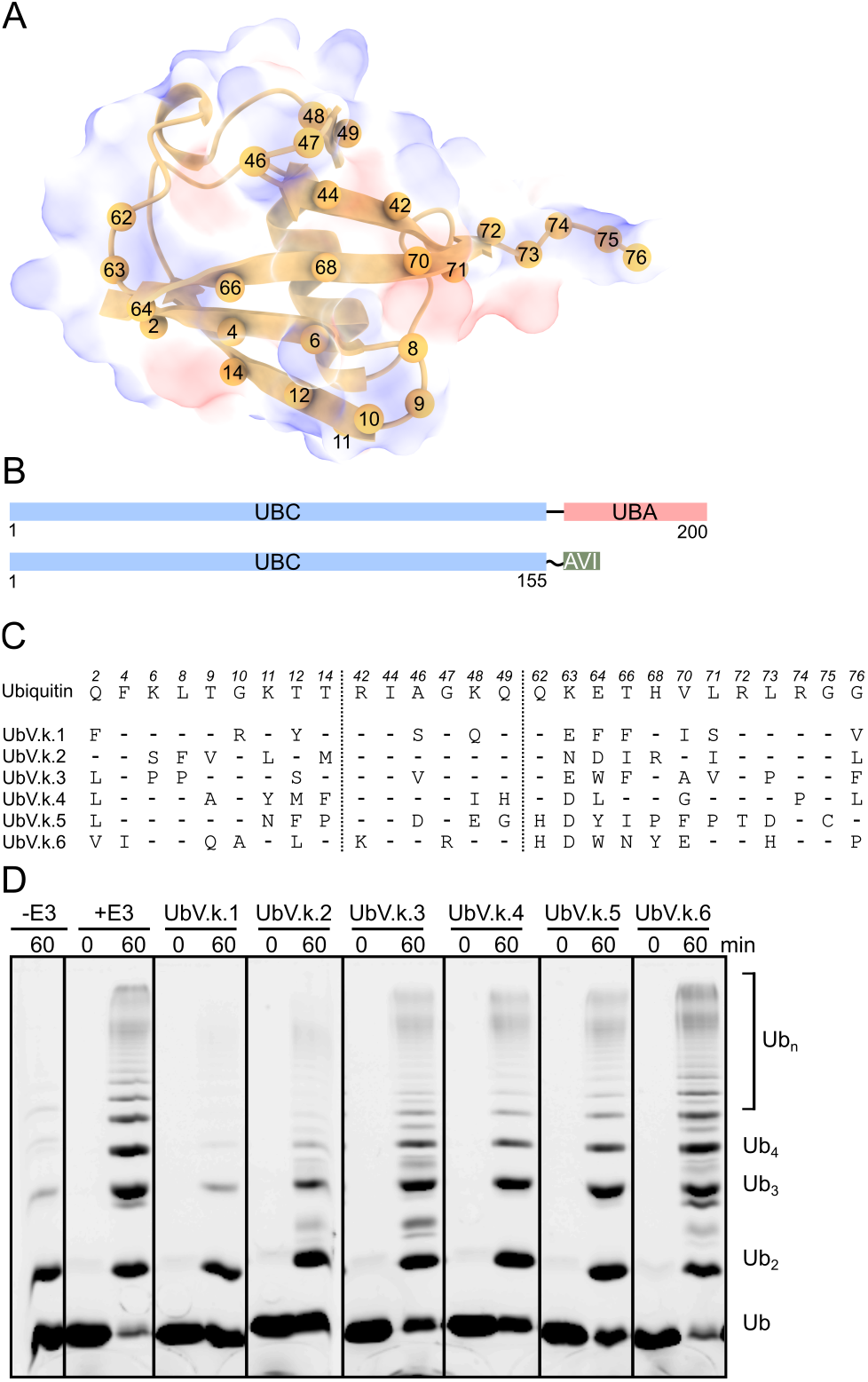
Selection for UbVs that bind to Ube2k. **(A)** The structure of wild-type ubiquitin (1UBQ) with the diversified residues shown as numbered spheres. Ubiquitin shown as a cartoon against a semi-transparent surface with acidic, basic, and hydrophobic groups coloured red, blue, or white, respectively. **(B)** Domain structure of full-length Ube2k and Ube2k-UBC used in this study. UBC: ubiquitin conjugating domain; UBA: ubiquitin associated domain; AVI: biotin-specific tag. **(C)** Sequence alignment of wild-type ubiquitin and the UbVs isolated from the selection. Only regions diversified in the library are shown. In the alignment, the amino acids indicate changes relative to the wild-type ubiquitin sequence, while dashes represent no change. **(D)** Chain-building ubiquitin (Ub) transfer assay performed without (-E3 and +E3) and with the six UbVs. For this assay, the E3 ligase RNF125 was used to promote ubiquitin transfer. Imaged as fluorescence from 5AIF-tagged ubiquitin. Assay performed in triplicate with similar results. See also Figure S1.

The Ube2k∼Ub conjugate was used for the screen in an attempt to isolate UbVs that could bind to either Ube2k, ubiquitin, or the E2-ubiquitin interface. Ube2k has a C-terminal extension that contains a ubiquitin associating (UBA) domain (Merkley and Shaw, 2004), which binds ubiquitin but is dispensable for ubiquitin transfer *in vitro*. As our goal was to identify modulators of ubiquitin transfer, we generated a truncated construct that comprised only the UBC domain of Ube2k (Ube2k-UBC). In addition, we included an active site mutation (C92K) to enable formation of a stable Ube2k∼Ub conjugate, and a C-terminal AVI tag to enable specific biotinylation of Ube2k (Figure 1B). After purification and biotinylation of Ube2k, the E2 enzyme was charged with ubiquitin, further purified, then immobilised to streptavidin/neutravidin-coated wells for phage-display selections.

After five rounds of phage-display selections against Ube2k∼Ub, clones were sequenced and six unique UbVs were chosen for closer analysis (Figure 1C). Each UbV (including an N-terminal FLAG tag) was subcloned into a vector that encoded a cleavable N-terminal His tag, and the resulting UbVs were expressed in *Escherchia coli*. After purification, interaction of each UbV with Ube2k∼Ub was confirmed using an ELISA experiment (Figure S1). Importantly, the ELISAs showed that each UbV bound to Ube2k and Ube2k∼Ub with comparable affinity, but did not bind ubiquitin alone. This indicated that the UbVs targeted Ube2k. The effect of each UbV on ubiquitin transfer was then assessed using a chain-building assay in the presence of Ube2k (Figure 1D). For this assay, each component of the ubiquitin cascade (E1, E2, E3, ubiquitin, and ATP) was mixed together and incubated at 37 °C prior to analysis. In this experiment, five of the UbVs (UbV.k.1-5 Figure 1C, 1D) decreased ubiquitin chain assembly by Ube2k, with UbV.k.1 and UbV.k.2 being the most potent inhibitors. As a result, we focussed our attention on these two UbVs.

### The inhibitory UbVs bind tightly and specifically to Ube2k

The UBC domain of E2 enzymes is structurally conserved, and while there is less conservation on a sequence level, there is the possibility that the UbVs that bind and inhibit Ube2k might cross-react with other E2s. To measure specificity, we assessed binding of UbV.k.1 and UbV.k.2 to a representative panel of 18 E2 enzymes. Our results revealed a preferential binding of both UbVs for Ube2k, with minimal binding to other E2 enzymes (Figure 2A). In support of this, no inhibition was seen when chain building assays were performed with two other E2 enzymes, Ube2d2 and Ube2n/Ube2v2 in the presence of the UbVs (Figure S2A).

**Figure 2.**
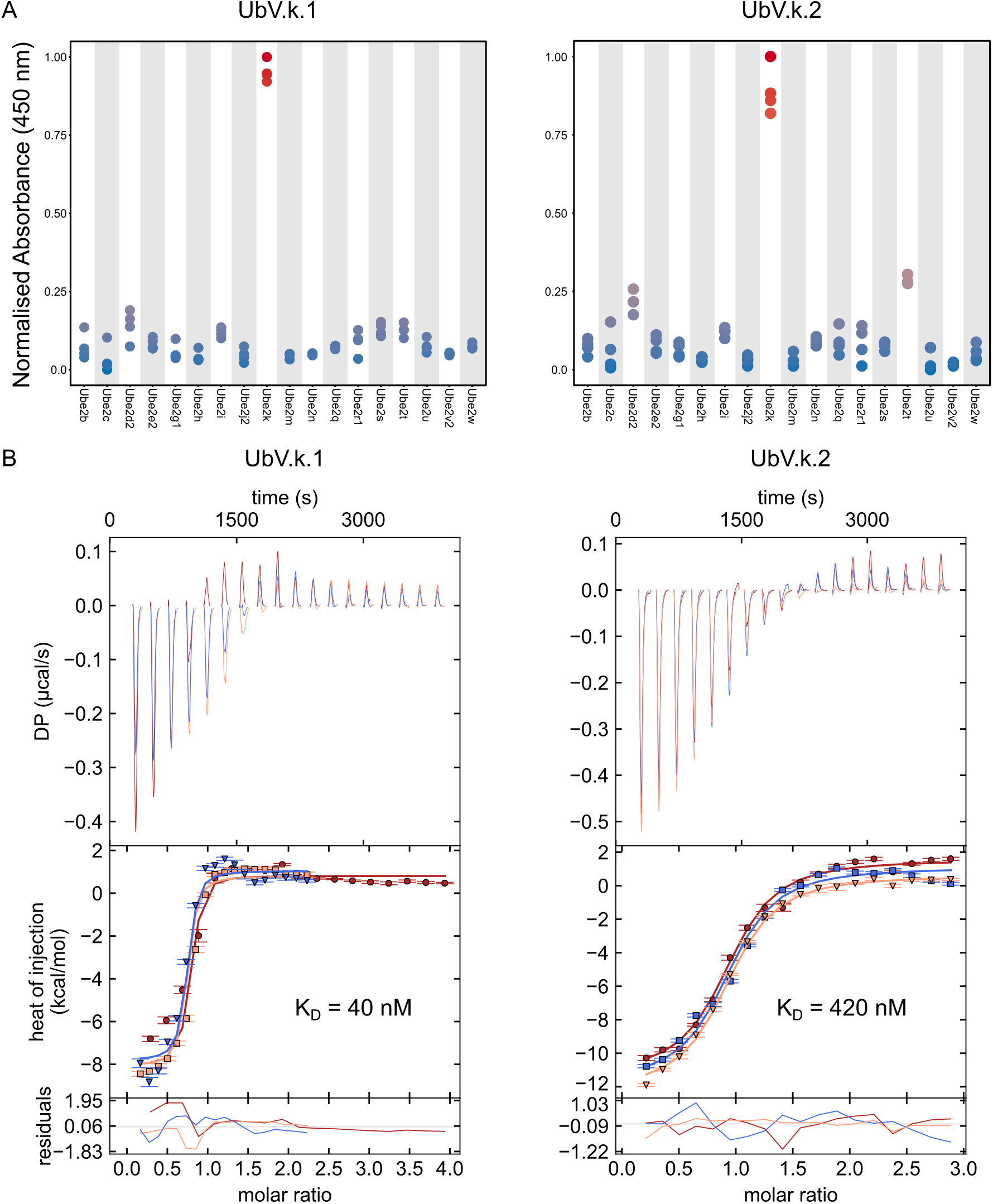
The UbVs bind to Ube2k specifically and tightly. **(A)** E2 enzyme specificity screen was performed using ELISA. A representative panel of E2 enzymes were immobilised to a plate before the UbVs were added and binding was detected spectrophotometrically by absorbance at 450 nm. Readings were normalised, and four replicates were performed. **(B)** Thermograms (top) and fitted isotherms (bottom) of ITC measurements performed in technical triplicate (shown in blue, red, and orange) where either UbV.k.1 or UbV.k.2 was injected into Ube2k. Binding constants (K_D_) are shown. See also Figure S2.

Next, we performed ITC to quantitatively measure the interaction between the UbVs and Ube2k. Each UbV was titrated against full-length Ube2k as well as Ube2k-UBC. The K_D_ of UbV.k.1 for Ube2k was calculated to be 40 nM (Figure 2B), while UbV.k.2 bound to Ube2k approximately ten-fold weaker, with a calculated K_D_ of 420 nM (Figure 2B). Similar dissociation constants were observed for both UbVs with Ube2k-UBC (Figure S2B, S2C), suggesting that the UbVs interacted with the UBC domain and not the UBA domain. In further support of a stable interaction, thermal denaturation experiments demonstrated that the melting temperature of Ube2k was increased by 4 or 2 °C in the presence of UbV.k.1 or UbV.k.2, respectively (Figure S2D). Together, these results demonstrate that the inhibitory UbVs are highly specific for Ube2k, bind tightly to the E2 enzyme, and increase its stability.

### The UbVs inhibit charging and discharging of Ube2k

What remained unclear was how the UbVs can block ubiquitin transfer from Ube2k. To promote ubiquitin transfer, E2 enzymes must first be loaded with ubiquitin by an E1 enzyme, which transfers ubiquitin to the E2 in a transthiolation reaction. The E2∼Ub conjugate then disengages from the E1 enzyme and binds an E3 ligase that catalyses ubiquitin transfer (Rennie et al., 2020). The chain-building assay (Figure 1D) involves both charging of the E2 by an E1 enzyme, and E3-dependent ubiquitin transfer. To tease out the mechanism of inhibition, we analysed E1-dependent charging and E3-catalysed transfer separately. First, to determine if the UbVs affected charging of Ube2k, we monitored the formation of an E2∼Ub thioester linked conjugate in the absence or presence of the UbVs. As observed in Figure 3A, both UbVs impeded charging of Ube2k as judged by slowed formation of an E2∼Ub species over time, with UbV.k.1 being a more potent inhibitor than UbV.k.2 (Figure 3A). We also confirmed that the UbVs did not affect the upstream E1 activation of ubiquitin (Figure S3). These results suggest that the UbVs impede loading of Ube2k by the E1 enzyme.

**Figure 3.**
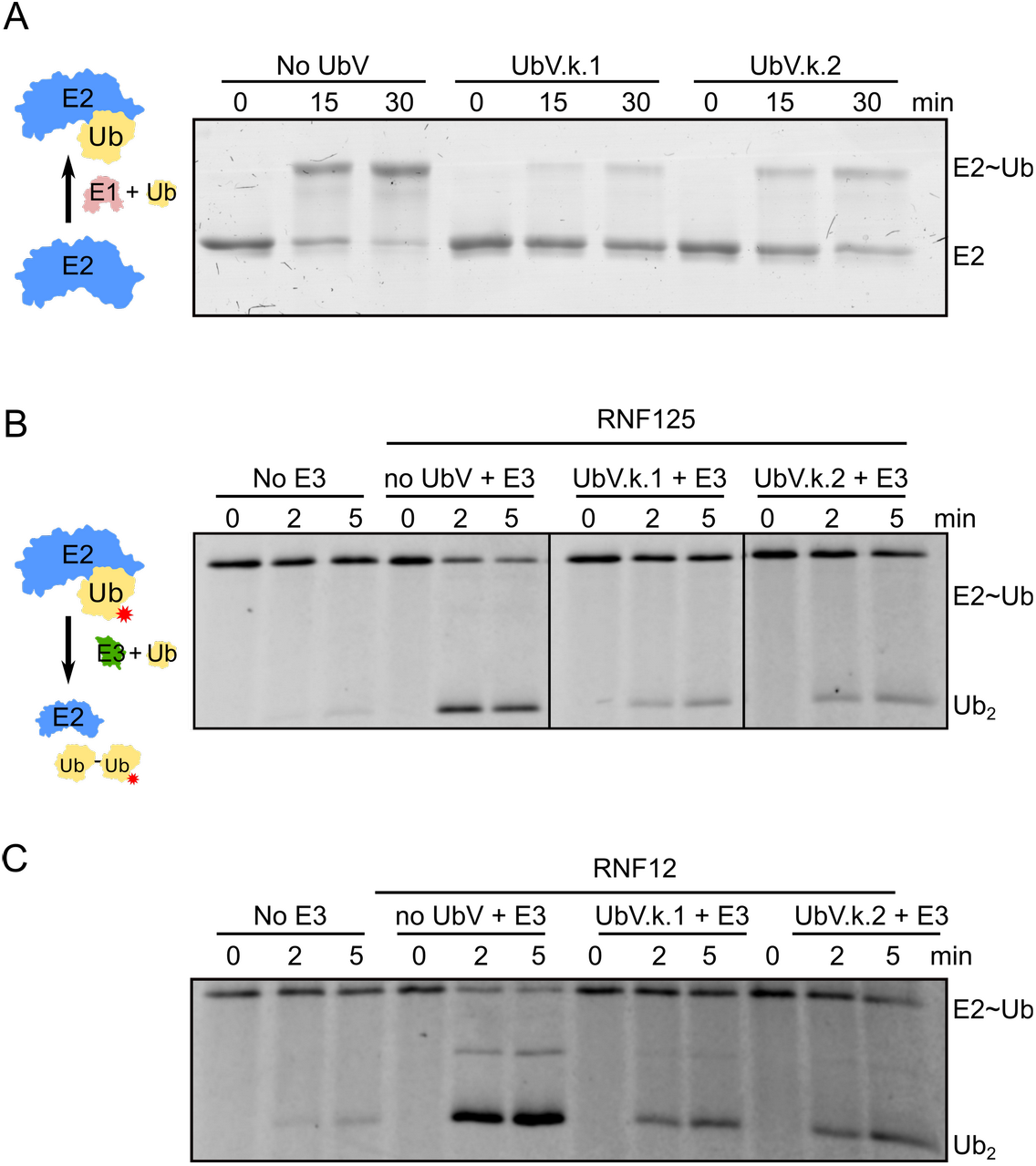
The UbVs inhibit both E2 charging and E3-catalysed ubiquitin discharge. **(A)** E2 charging assay (see schematic on the left) performed with and without the UbVs. Samples were quenched with non-reducing dye at the indicated time points. The gel was visualised by staining with Coomassie blue dye. **(B, C)** A ubiquitin discharge experiment (schematic on the left; red star indicates fluorescent tag) performed with and without the UbVs. Samples were quenched with non-reducing dye at the indicated time points. The gel was visualised by fluorescence detected at 600 nm. Panels (B) and (C) show results with RNF125 or RNF12 as the E3 ligase, respectively. See also Figure S3 and S4.

To assess whether the UbVs could also inhibit E3-catalysed ubiquitin transfer from Ube2k, we prepared a fluorescently-tagged thioester-linked Ube2k∼Ub-Cy3^K0^ conjugate and monitored the disappearance of the conjugate and appearance of di-ubiquitin in the presence of an E3 ligase and excess UbVs (Figure 3B). Our results show that the E3-dependent discharge was slowed by the UbVs when using either RNF12 or RNF125 E3 ligases (Figure 3B, 3C). However, in the absence of an E3 ligase, the UbVs did not affect basal ubiquitin transfer activity of Ube2k (Figure S4). These results suggest that, as well as inhibiting charging of the E2 enzymes by the E1 enzyme, the UbVs also disrupted E3 ligase catalysed ubiquitin discharge.

### Crystal structures of the UbV-Ube2k complexes reveal the mechanism of inhibition

To establish exactly how the two UbVs bind and inhibit the activity of Ube2k, we solved their structures in complex with the E2 enzyme. Following copurification of UbV.k.2 and Ube2k, crystals were obtained for the complex and its structure was solved by molecular replacement using Ube2k and ubiquitin. The structure was refined to a resolution of 2.4 Å with final R_work_/R_free_ of 20.6 / 24.9 (Table 1). Crystals of a similarly purified UbV.k.1-Ube2k complex could not be obtained. Instead, we purified an isopeptide-linked Ube2k∼Ub conjugate and copurified it with UbV.k.1. With this complex, crystals were obtained and the structure of UbV.k.1-Ube2k∼Ub was solved to 3.0 Å with final R_work_/R_free_ of 23.4 / 29.3 (Table 1). In both cases, the asymmetric unit contained only one complex.

In both structures, Ube2k shows the expected fold comprising a UBC domain bridged by a short linker to a C-terminal UBA domain (Figure S5A). A C-alpha overlay of Ube2k from the two structures has an RMSD of 1.2 Å, and they both overlay well with a published structure of Ube2k (PDB: 5DFL)(Middleton and Day, 2015) with C-alpha RMSDs of 1.3 and 0.65 Å for Ube2k-UbV.k.1 and -UbV.k.2, respectively. In the Ube2k-UbV.k.2 complex, residues 32-34 of Ube2k and 7-12 of UbV.k.2 are shifted towards each other, and it appears that a slight conformational change of both proteins is necessary to form the complex.

Both UbVs bind at the same position of Ube2k, which comprises a hydrophobic cleft formed between α-helix 1 and β-sheet 1 of Ube2k (Figure 4A, 4B). Surprisingly, an overlay of the two complexes shows that the UbVs are rotated by approximately 15° with respect to one another and Ube2k (S5B), even though the contact residues from both are largely from the same sequence positions. Notably, the contact residues from both UbVs are part of the diversified surface, with major contributions from those mutated from wild-type ubiquitin (Figure 1B, Figure 4C, 4D, 4E). An analysis using PISA (Protein, Interfaces, Structures and Assemblies) (Krissinel and Henrick, 2007) reveals that the interface between UbV.k.1 and Ube2k buries approximately 470 Å^2^, while UbV.k.2 buries approximately 530 Å^2^. While UbV.k.2 buries a greater surface area of Ube2k than UbV.k.1, the smaller hydrophobic patch at the core of the UbV.k.2-Ube2k interaction may explain its reduced affinity for Ube2k (Figure 4E, 4F).

**Figure 4.**
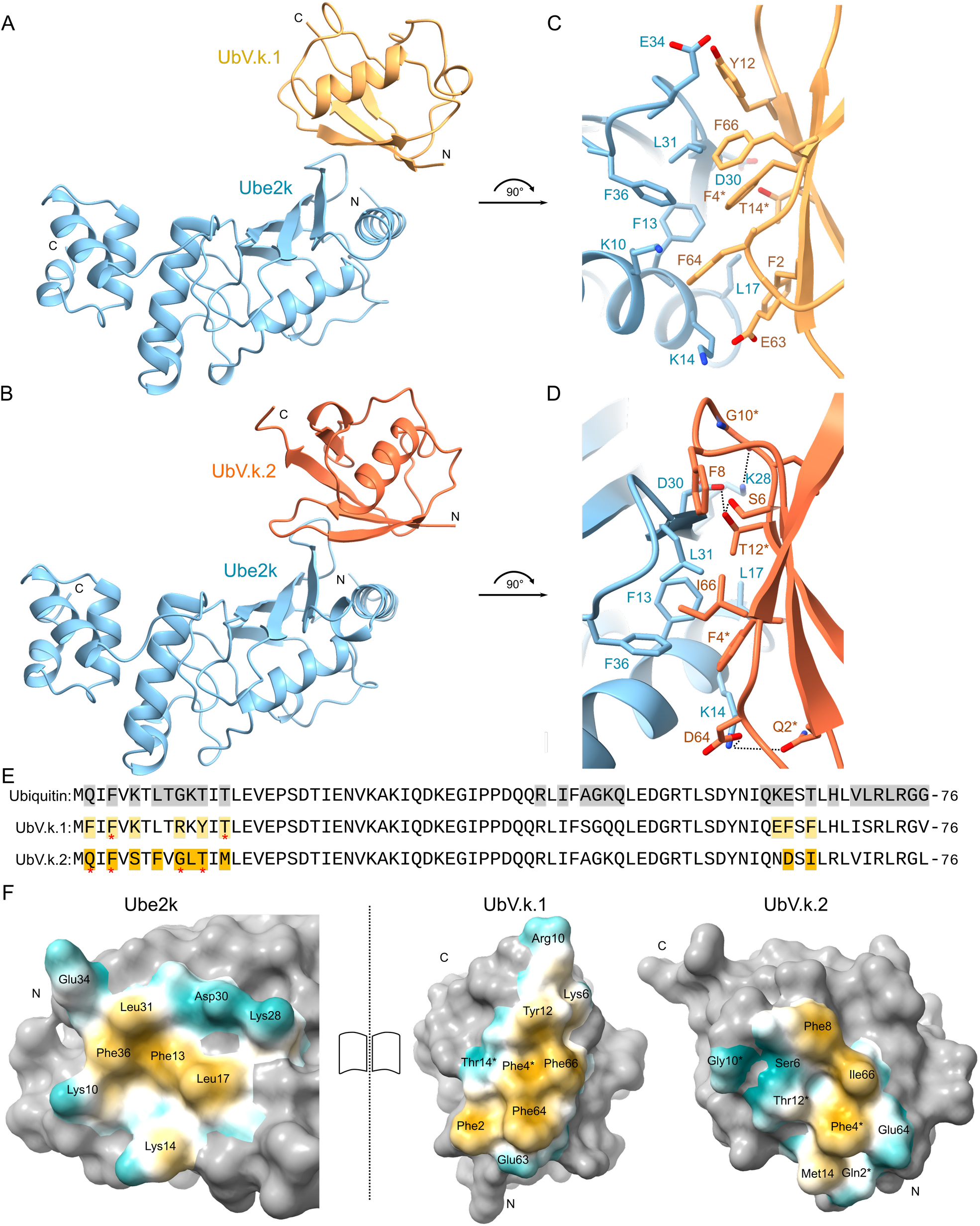
Crystal structures of UbV.k.1 or UbV.k.2 in complex with Ube2k. **(A,B)** Ribbon representation of UbV.k.1 (A) or UbV.k.2 (B) in complex with Ube2k. N and C termini are labelled. **(C)** Sequence alignment of wild-type ubiquitin (Wt-Ub), UbV.k.1 and UbV.k.2 with the residues that contact Ube2k highlighted. Red asterisks indicate residues that are unchanged from wild-type ubiquitin. **(D)** Open book representation of the UbV-Ube2k interactions. On the left is Ube2k from both complexes, while UbV.k.1, and UbV.k.2 are shown on the right. Non-contact residues are shown in grey, while contact residues are coloured based on their molecular lipophilicity potential: hydrophobic atoms are coloured orange, while hydrophilic atoms are cyan. Residues unchanged from wild-type ubiquitin are indicated with asterisks. **(E**,**F)** Close up views of the interfaces between the UbVs and Ube2k. The main chain are shown as ribbons and side chains are shown as sticks coloured as in panels (A) and (B). Dashed lines indicate predicted hydrogen bonds. Residues unchanged from wild-type ubiquitin are indicated with asterisks. See also Figure S5.

The more potent UbV, UbV.k.1, uses an array of four Phe residues (Phe2, Phe4*, Phe64, and Phe66; wild-type ubiquitin residues are indicated with asterisks) and one Tyr residue (Tyr12) to form a hydrophobic surface that packs against Ube2k residues Phe13, Leu17, Leu31, and Phe36 (Figure 4C, 4F). This patch is complemented by a hydrophilic ‘shell’ made up of the hydroxyls of Tyr12, Thr14*, and the side chain Glu63 of UbV.k.1; and Lys10, Lys14, Asp30, and Glu34 of Ube2k (Figure 4C, 4F). In a similar manner, the UbV.k.2-Ube2k interaction has a hydrophobic core, comprising Phe4*, Phe8, and Ile66 of UbV.k.2, which interacts with the same patch on Ube2k as for UbV.k.1 (Phe13, Leu17, Leu31, and Phe36). This is supported by an extensive polar shell nucleated by Ser6 of UbV.k.2, which acts to stabilise contacts between Thr12* of UbV.k.2 with Asp30 and Leu31 of Ube2k (Figure 4D, 4F). In addition, the side chain of Lys14 of Ube2k contacts Gln2* and Asp64 of UbV.k.2, while the side chain of Lys28 of Ube2k makes contacts with Gly10*.

To test the importance of the diversified residues, we introduced single point mutations to revert both UbVs to the wild-type ubiquitin sequence and assessed binding of the mutants to Ube2k by ELISA EC_50_ measurements. Interestingly, each revertant disrupted the EC_50_ by at least an order of magnitude, while some of the mutations almost eliminated binding (Figure 5A, 5B). For UbV.k.1, mutation of Phe64 and Glu63 to their wild-type amino acid (Glu and Lys, respectively) almost completely disrupted binding to Ube2k, while for UbV.k.2, mutating Ser6 to Lys, as well as Ile66 to Thr were highly disruptive to binding. In support of the importance of these residues, the mutations, F64E and E63K made to UbV.k.1, as well as S6K and I66T on UbV.k.2, did not inhibit the formation of ubiquitin chains (Figure 5C). These data suggest that the network of diversified residues is highly interdependent and disruption of any of them results in a notable loss of binding and inhibition.

**Figure 5.**
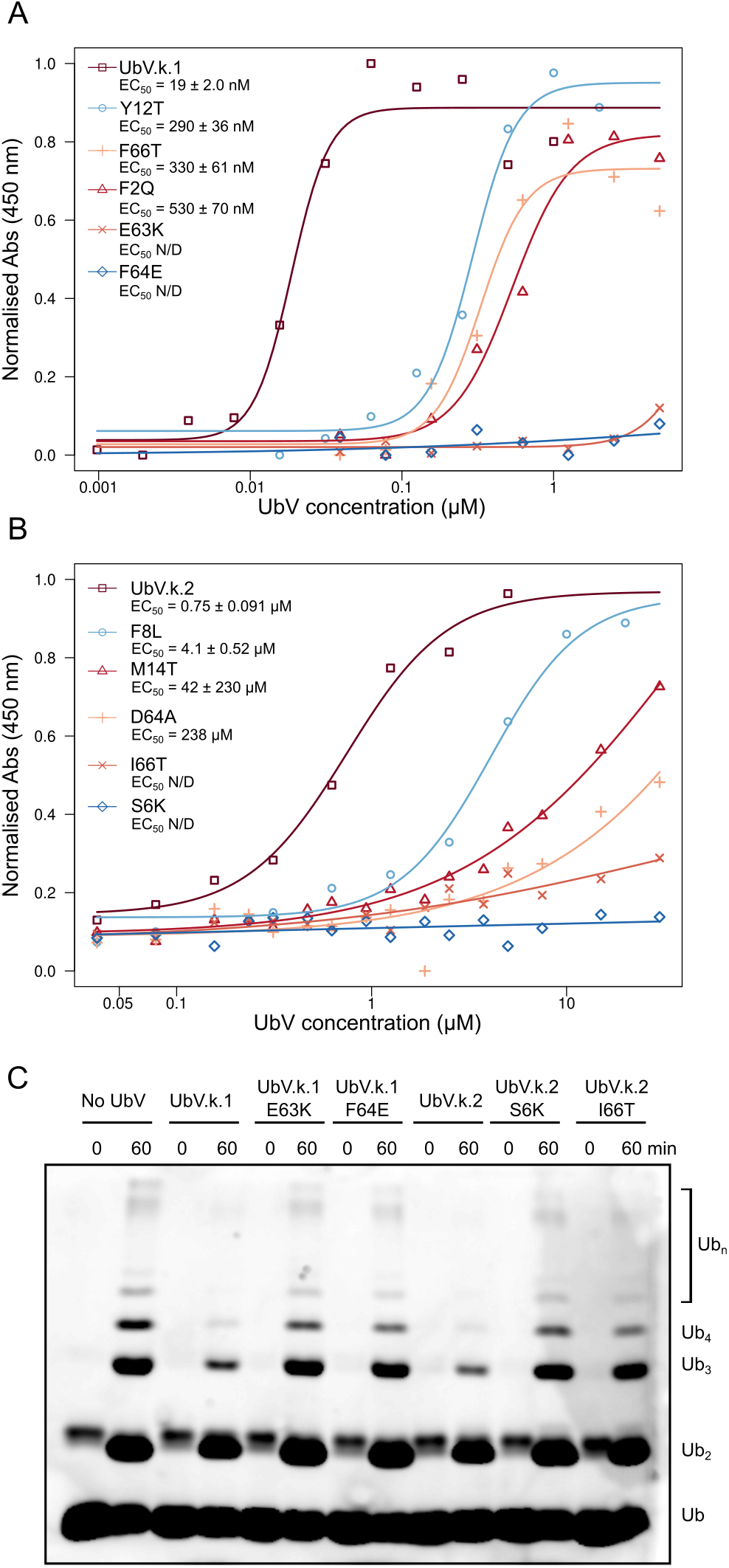
Single point mutations to UbVs are sufficient to decrease both Ube2k binding and inhibitory activity. **(A,B)** The binding of the UbVs and their mutants to Ube2k was detected using an ELISA as in Figure 2A. UbV.k.1 and its mutants are shown in panel (A), while UbV.k.2 and its mutants are in panel B. The EC_50_ values were calculated as the point where the absorption signal was 50% of maximum. **(C)** Ubiquitin chain-building assay performed with or without the UbVs or revertants that fully disrupted binding. Imaged as fluorescence from 5AIF-tagged ubiquitin.

### Architecture of UbV-Ube2k interactions

Ubiquitin activation by an E1 enzyme sets the stage for subsequent transfer to E2 enzymes and E3-ligase catalysed ubiquitin transfer. The E1 enzyme engages E2s using two major interactions: first the E2 enzyme is recruited by a ubiquitin-fold domain (UFD) that interacts with a semi-conserved surface on E2s; subsequently, the UFD domain rotates and positions the active site domain of the E1 next to the catalytic Cys of the E2 enzyme to allow a transthiolation reaction to occur (Olsen and Lima, 2013; Williams et al., 2019). Because there is no structure of Ube2k bound to the E1, we generated a molecular model by overlaying Ube2k from our two structures with Cdc23 from a recent E2-E1 complex structure and mapped the predicted UFD contacts on Ube2k (Figure 6A). In our model, both the UbVs clash with the predicted binding site of the UFD, likely explaining why ubiquitin charging of Ube2k by the E1 enzyme is reduced in the presence of the inhibitory UbVs. The UbVs also disrupt the activity of E3-catalysed discharge of Ube2k (Figure 3B, 3C). To assess whether the UbVs clash with the likely E3 binding site, we generated a model of an E3-engaged Ube2k molecule. For this, we overlayed Ube2k from both complexes with Ube2d2 from a crystal structure of Ube2d2-RNF12 (Figure 6B) (Middleton et al., 2020). In this model there is no overlap between RNF12 and the UbVs, suggesting that inhibition of E3-catalysed discharge of Ube2k is likely due to allosteric effects. Together, the architecture of both the UbV-Ube2k interactions suggests a direct steric clash with the E1 enzyme, while the mechanism of inhibition of the E3 ligase by the UbVs is less clear, and appears to be due to allosteric effects.

**Figure 6.**
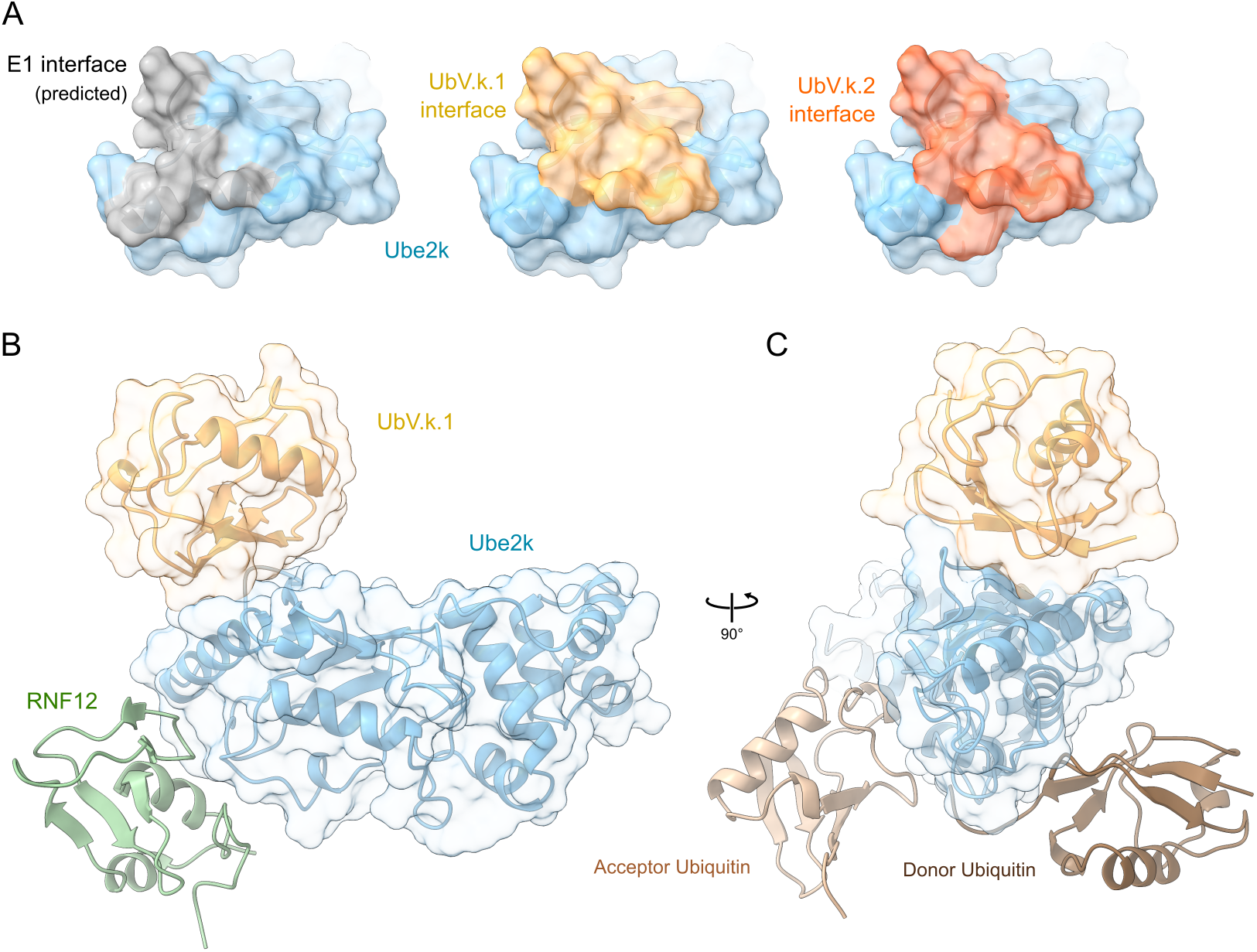
Molecular architecture of the Ube2k-UbV structures. **(A)** Surface representation of Ube2k showing residues predicted to be within 3 Å of the UFD domain of the E1 ubiquitin activating enzyme (left, in grey), and contacts with UbV.k.1 or UbV.k.2 (middle and right in yellow or orange, respectively). Non-contacting Ube2k residues are shown in blue. **(B)** Model of RNF12-bound Ube2k was generated by overlaying the E2 molecules from the UbV.k.1-Ube2k and UbV.k.2-Ube2k structures with Ube2d2 from the RNF12-Ube2d2 crystal structure (PDB ID: 6W7Z). Only one Ube2k molecule with UbV.k.1 is shown represented in ribbon and semi-transparent surface. The modelled RNF12 is shown as a green ribbon. The dotted line shows the measured distance between RNF12 and the UbVs. **(C)** Ube2k in complex with UbV.k.1 with donor and acceptor ubiquitin molecules modelled at their predicted sites (Middleton and Day, 2015).

## Discussion

As the central enzymes in the ubiquitin cascade, E2 enzymes play a critical role in determining the exact nature of the ubiquitin code (Stewart et al., 2016). Here we have focused on developing tools to modulate targeting of proteins to the proteasome by identifying regulators of the E2 enzyme, Ube2k, which only produces Lys48-linked degradative ubiquitin chains. Using phage display, we discovered two UbVs that are potent inhibitors of ubiquitin transfer by Ube2k. The UbVs bind tightly and highly specifically to Ube2k, and they appear to isolate the E2 from the ubiquitin system by disrupting interactions with both E3 ligases and the E1 enzyme. The fact that both UbVs bind at the same hydrophobic cleft on Ube2k suggests this site may prove to be an effective target for small molecule binding. By inhibiting the synthesis of a degradative signal, these UbVs will provide tools for research, and may suggest approaches for design of small molecule Ube2k inhibitors. For example, the UbVs may prove useful for discovery of small-molecules that bind at this same hydrophobic cleft by displacement assays, as demonstrated by others (Maculins et al., 2020).

To date, approximately twenty inhibitors of ubiquitin-conjugating E2 enzymes have been discovered (Ardecky et al., 2010; Ceccarelli et al., 2011; Chen et al., 2017; Cheng et al., 2014; Cornwell et al., 2019; Garg et al., 2020; Helms et al., 2011; Kothayer et al., 2016; Liu et al., 2014; Pulvino et al., 2012; Sanders et al., 2013; Strickson et al., 2013; Wang et al., 2018, 2011). Many of the reported inhibitors target the active site Cys, others act by disrupting protein-protein interactions, and some have an allosteric effect on ubiquitin transfer. Because of the conservation and lack of a distinctive active site pocket on E2 enzymes, many of these orthosteric inhibitors have low specificity and relatively high IC_50_ values (Osborne et al., 2021). Display technologies offer powerful ways of isolating protein or peptide-based inhibitors of proteins (including E2 enzymes) that can circumvent these problems. For example, phage display was used to discover UbVs that bound at the ‘backside’ ubiquitin binding site found on some E2 enzymes (Garg et al., 2020). These variants bound more tightly than many other inhibitors of E2 enzymes, with IC_50_ measurements ranging from 65-280 nM. Importantly, they could all inhibit ubiquitin transfer by either directly blocking interactions with other components of the ubiquitin cascade (similar to what we observed here) or by allosteric effects that disrupt the enhanced ubiquitin transfer normally provided by backside-bound ubiquitin.

Surprisingly, the UbVs we discovered did not bind at a predicted ubiquitin-binding site (Figure 6C), nor did they block the active site Cys. Instead they bound at a hydrophobic groove between α-helix 1 and β-sheet 1 on Ube2k that overlaps considerably with the binding site of the UFD domain of E1 enzymes (Figure 6A) (Lv et al., 2017; Olsen and Lima, 2013; Tokgoz et al., 2011; Williams et al., 2019), but binds in a distinct manner. The UFD-E2 interface appears to be largely governed by polar contacts between acidic residues on the E1 and a set of basic residues conserved on helix-1 of E2 enzymes. By contrast, both of the UbVs reported here rely on a hydrophobic patch, as well as critical polar contacts between the molecules (Figure 4C, 4D, 4F, Figure 5). A sequence comparison of the E2 enzymes used in our representative screen suggests that the residues of Ube2k that interact with the UbVs are not highly conserved (Figure S6). In particular, this is true for the hydrophobic residues, Phe13, Leu17, Leu31, and Phe36, which provide the core of both of the interactions. This low conservation likely explains the specificity of the interaction with Ube2k. How the UbVs block engagement by the E3 ligase is not as clear, because in our model (Figure 6B) the UbVs bind away from the predicted RNF12 binding site on Ube2k. E3 ligase-E2 interactions are highly conserved, and it is unlikely that the E3-Ube2k interaction will differ greatly from our model. As a result, it is probable that the inhibition of E3 ligase activity is due to allosteric effects caused by the UbV-Ube2k interactions.

Currently, the role of Ube2k in biology has not been extensively investigated, but studies have shown that the enzyme appears to have diverse roles in cells, such as regulating cell division (Bae et al., 2013), controlling the stability of p53 (Hong et al., 2019), and is likely to play roles in the development of neurodegenerative diseases (Kalchman et al., 1996; Meiklejohn et al., 2019; de Pril et al., 2007; Song et al., 2003; Su et al., 2018). The UbVs may prove to be productive in narrowing down the exact importance of Ube2k in the cell. While there is considerable redundancy in the cell, Ube2k is the only E2 enzyme that contains a ubiquitin-binding domain, which may act to increase the local concentration of the enzyme to promote rapid synthesis of extensive ubiquitin chains, and degradation. An interesting recent report suggests the UBA domain may be involved in recruiting Ube2k to Lys63-linked ubiquitin chains to potentially quench signalling pathways promoted by Lys63 chains (Pluska et al., 2021). Further work is needed to understand more about the role of Ube2k in cells, and the UbVs may prove essential to these studies.

## Materials and Methods

### DNA Constructs

For phage selection, a DNA construct encoding residues 1-157 of Ube2k (Ube2k-UBC) containing the mutations C92K and K97R fused to a short GSGS linker and an AVI tag was obtained from GeneArt (Invitrogen). This construct was used to generate a stable isopeptide-linked Ube2k∼Ub conjugate. A DNA construct encoding the biotin ligase BirA was obtained from GeneArt. Both of these constructs were cloned into a pET-LIC vector encoding a 6xHis tag followed by a 3C protease cleavage site N-terminal to the construct. For the E2 specificity screen, a modified pET-LIC vector was constructed to allow co-expression of C-terminal AVI-tagged proteins with the biotin ligase BirA. This vector was named pET-LIC-3Avi-BirA-Duet. For cloning of the E2s, primers containing LIC overhangs were used to amplify a representative panel of E2 enzymes, and these were then cloned into pET-LIC-3Avi-BirA-Duet. Mutations were generated using single-step site-directed mutagenesis (Liu and Naismith, 2008). After phage selection, the DNA sequences encoding the UbVs were amplified using universal primers and cloned into pET-LIC. Cloned UbVs included an N-terminal FLAG tag and a C-terminal GGSGG tag. An RNF125 construct encoding residues 16-125, as well as Ube2k were cloned into pGEX-6P3 using BamHI and EcoRI restriction digestion and ligation.

### Protein expression and purification

Each protein was produced in the *E. coli* BL21-star cell line in LB broth by incubation at 37 °C followed by 18 °C for overnight expression. Protein expression was either induced by 0.2 mM IPTG, or by autoinduction. After centrifugation, cell pellets were resuspended in 1x PBS, sonicated, and centrifuged at 15,000x g. For constructs in the pET-LIC vector, the His-tagged protein was immobilised to nickel beads (Macherey-Nagel), eluted with 300 mM imidazole, and either dialysed or desalted into PBS. Subsequently, this was mixed with in-house produced His-tagged 3C for overnight digestion to remove the His tag. For purification of BirA, the 3C digestion was not performed. After confirmation of digestion, the protein was further purified using size-exclusion chromatography over a Superdex S75 13/300 or 16/600 column (GE Healthcare). For Ube2k-UBC containing the Avi tag, the purified protein was incubated at a final concentration of 100 μM with 300 μM biotin, 2 μM BirA, 2.5 mM ATP, and 2.5 mM MgCl_2_ for 1 h at room temperature. To confirm biotinylation, the purified proteins were incubated with excess streptavidin (New England Biolabs) for 10 minutes before analysis via gel-shift using SDS-PAGE. The protein was then desalted to remove excess biotin, and the BirA removed using Ni^2+^ affinity chromatography. For the constructs in pET-LIC-3Avi-BirA-Duet, a final concentration of 200 μM biotin was added to the *E. coli* cultures before transfer to 18 °C. Ubiquitin, E1, Ube2k, Ube2d2, Ube2n, Ube2v2, RNF12, TRAF6 and 3C protease were purified using previously established methods (Berndsen and Wolberger, 2011; Middleton and Day, 2015; Middleton et al., 2017, 2020; Sato et al., 2009). RNF125 was purified as described for RNF12. Ubiquitin with all Lys residues mutated to Arg (K0) was purified identically to wild-type ubiquitin. Thioester and isopeptide linked Ube2k∼Ub were purified using techniques previously described(Middleton et al., 2014). Fluorescently labelled ubiquitin for chain-building assays (Ub-5AIF), ubiquitin discharge assays (Ub-Cy3^K0^) and E1 activation assays (Ub-Cy3) were produced as previously described. For the pulldown experiment, GST-Ube2k and GST-3C were bound to GSH resin and washed extensively. GST-Ubiquitin and GST alone were purified by binding to GSH resin, and elution from the resin with 10 mM GSH for 30 min. Subsequently, they were purified by size-exclusion chromatography. All proteins were frozen in liquid nitrogen and kept at -80 °C.

### Phage-display selections

Five rounds of phage-display selections were performed as previously described using a library highly similar to the ones used in earlier studies (Ernst et al., 2013; Garg et al., 2020). For selection, biotinylated Ube2k-UBC∼Ub was immobilised to Nunc Maxisorp 96-well plates (Fisher Scientific, Nepean, ON, Canada) coated alternately with neutravidin (New England BioLabs) or streptavidin (ThermoFisher Scientific). After blocking, the UbV library was added to immobilised Ube2k∼Ub for 1 h at 4 °C followed by washing with PBS containing 0.02% Tween-20 and elution at low pH. Subsequently, *E. coli* TOP10 cells (ThermoFisher Scientific) were infected with the eluted phage before being grown overnight at 37 °C. The *E. coli* TOP10 cells were plated and 96 individual clones (48 from round four and 48 from round five) were grown and sequenced. From the sequences, six distinct UbVs were analysed in subsequent experiments.

### Ubiquitin transfer assays

For ubiquitin chain-building experiments, 5 μM E3 (RNF125, TRAF6, or RNF12), 0.1 μM E1, 8-10 μM E2 (Ube2k, Ube2d2, or Ube2n/Ube2v2), 50 μM ubiquitin, and 5 μM Ub-5AIF were incubated with or without 10-30 μM UbVs in buffer containing 20 mM Tris-HCl (pH 7.5), 50 mM NaCl, 2 mM TCEP, 2 mM ATP, and 2 mM MgCl_2_. Reactions were incubated at 37 °C and mixed with SDS loading dye containing 2-mercaptoethanol at the indicated time points. The resulting SDS-PAGE gels were imaged on a Las-3000 (Fuji Film) imager for approximately 60 s. For ubiquitin discharge assays, purified thioester-linked Ube2k∼Ub-Cy3^K0^ conjugate (approximately 10 μM) was mixed with 250 nM RNF125 or 2.5 μM RNF12 and 50 μM ubiquitin with or without UbVs, and incubated at 20 °C. At the indicated time points, the samples were mixed with SDS loading dye containing no reducing agent and resolved with SDS-PAGE. Samples were imaged on an Odyssey FC imaging system (LI-COR) at 600 nm with a 2 min exposure. Charging assays of Ube2k were performed in PBS with 0.1 μM E1, 10 μM Ube2k, 50 μM ubiquitin, +/- 10 μM UbV.k.1 or UbV.k.2, 2 mM MgCl_2_, and 2 mM ATP. Reactions were incubated at 37 °C before being mixed with non-reducing SDS dye and proteins were visualised with Coomassie blue staining of the gels. E1 activation experiments were performed in PBS, 2 mM MgCl_2_, 2 mM ATP with 0.5 μM E1, 50 μM Ub-Cy3, with or without 30 μM UbV.k.1 or UbV.k.2. The reactions were incubated at 20 °C for 5 min before being mixed with non-reducing SDS dye, and gels resolved via fluorescence of Cy3 as for discharge experiments.

### ELISA experiments

For epitope mapping, 384-well high-binding plates (Greiner Bio-One) were coated overnight with streptavidin, GST, or GST-Ubiquitin. Biotinylated Ube2k-UBC∼Ub, Ube2k, or PBS was mixed with the immobilised streptavidin for ∼ 15 min. Purified UbVs were then added at 0.5 μM and binding was detected via reaction of 3,3’,5,5’-tetramethylbenzidine (TMB) with an HRP-FLAG antibody (1:4000, ThermoFisher Scientific). The colorimetric reaction was quenched with either phosphoric or sulfuric acid after 5-10 min. Absorbance at 450 nm was measured with a ClarioSTAR Plus (BMG Labtech). Background was subtracted and values were normalised where the highest signal was 1.0. Similar approaches were used for the E2 specificity and EC_50_ ELISA experiments. For the EC_50_ measurements, the UbVs were diluted as indicated before being mixed with immobilised Ube2k. The EC_50_ value is the concentration at which the signal of absorbance is 50% of total binding. These data were analysed and plotted in R version 4.0.4.

### Binding experiments

Isothermal titration calorimetry was performed with a VP-ITC (MicroCal) at 30 °C. Ube2k and Ube2k-UBC were added to the cell at 8 and 8-10 μM, respectively, while UbV.k.1 (at 150 or 85 μM) and UbV.k.2 (at 110 μM) were in the syringe. All samples were either dialysed against or purified with a common stock of PBS. Analysis was performed using NITPIC, SEDPHAT, and GUSSI (Scheuermann and Brautigam, 2015). Thermal denaturation of Ube2k was performed at a final protein concentration of 5 μM. SYPRO Orange (ThermoFischer) dye was added to the protein in a white 96-well PCR plate (Lab Supply, Dunedin, New Zealand) and measured in a Roche LightCycler 480 II instrument using the SYPRO Orange program. Data were analysed and plotted with R version 4.0.4.

### Crystallography and structure solution

For crystallography, the copurified complexes were mixed with the crystal screens PACT Premier and JCSG plus (Molecular Dimensions) at 200:200 nL and 200:100 nL protein:well solution drop ratios in Swissci 3-well sitting drop plates using a mosquito (TTP Labtech). Diffraction-quality crystals of the UbV.k.2-Ube2k complex were produced in 0.2 M sodium citrate tribasic trihydrate, 0.1 M Bis-Tris propane pH 7.5, and 20% PEG 3350 and the dataset was collected at the MX1 beamline, Australian Synchrotron. UbV.k.1-Ube2k∼Ub crystals were grown in 0.2-0.3 M ammonium citrate dibasic, 20-25% PEG 3350 and data were collected at the MX2 beamline, Australian Synchrotron. Data were processed and scaled with XDS (Kabsch, 2010), and datasets were merged with Aimless (CCP4, 1994). Subsequently, Phaser-MR (McCoy et al., 2007) was used to solve the structures using a single Ube2k molecule from PDB 5DFL (Middleton and Day, 2015), and the core domain of ubiquitin (PDB: 1UBQ) (Vijay-Kumar et al., 1987). For the UbV.k.1-Ube2k∼Ub dataset, the ice ring at 3.4 Å was excluded from the processing. Both structures were refined with Phenix refine (Adams et al., 2010), and manually corrected iteratively using Coot (Emsley et al., 2010). All images were generated using ChimeraX (Pettersen et al., 2021). Crystal coordinates for UbV.k.1-Ube2k∼Ub and UbV.k.2-Ube2k have been deposited in the Protein Data Base under accession codes 7MYF or 7MYH, respectively.

## Acknowledgements

We acknowledge Jun Gu for assistance with phage display, and Khanh Nguyen for help with the thermal denaturation experiments. This research was undertaken in part using the MX1 and MX2 beamline at the Australian Synchrotron, part of ANSTO, and made use of the Australian Cancer Research Foundation (ACRF) detector. We thank the New Zealand Synchrotron Group for facilitating access to the Australian Synchrotron. A.J.M. was supported by a Marsden Fund Fast-start Grant administered by the Royal Society of New Zealand, and a Sir Charles Hercus Health Research Fellowship awarded by the Health Research Council of New Zealand.

## Author Contributions

Conceptualization, A.J.M., C.L.D., S.S.S.; Investigation, A.J.M., J.Z., J.T.; Resources, A.J.M., S.S.S., C.L.D.; Writing – Original Draft, A.J.M.; Writing – Review & Editing, C.L.D., S.S.S., J.T., A.J.M.; Visualization, A.J.M., J.T.; Funding Acquisition, A.J.M., S.S.S., C.L.D.

## Declaration of Interests

The authors declare no competing interests.

---

**Figure S1.**
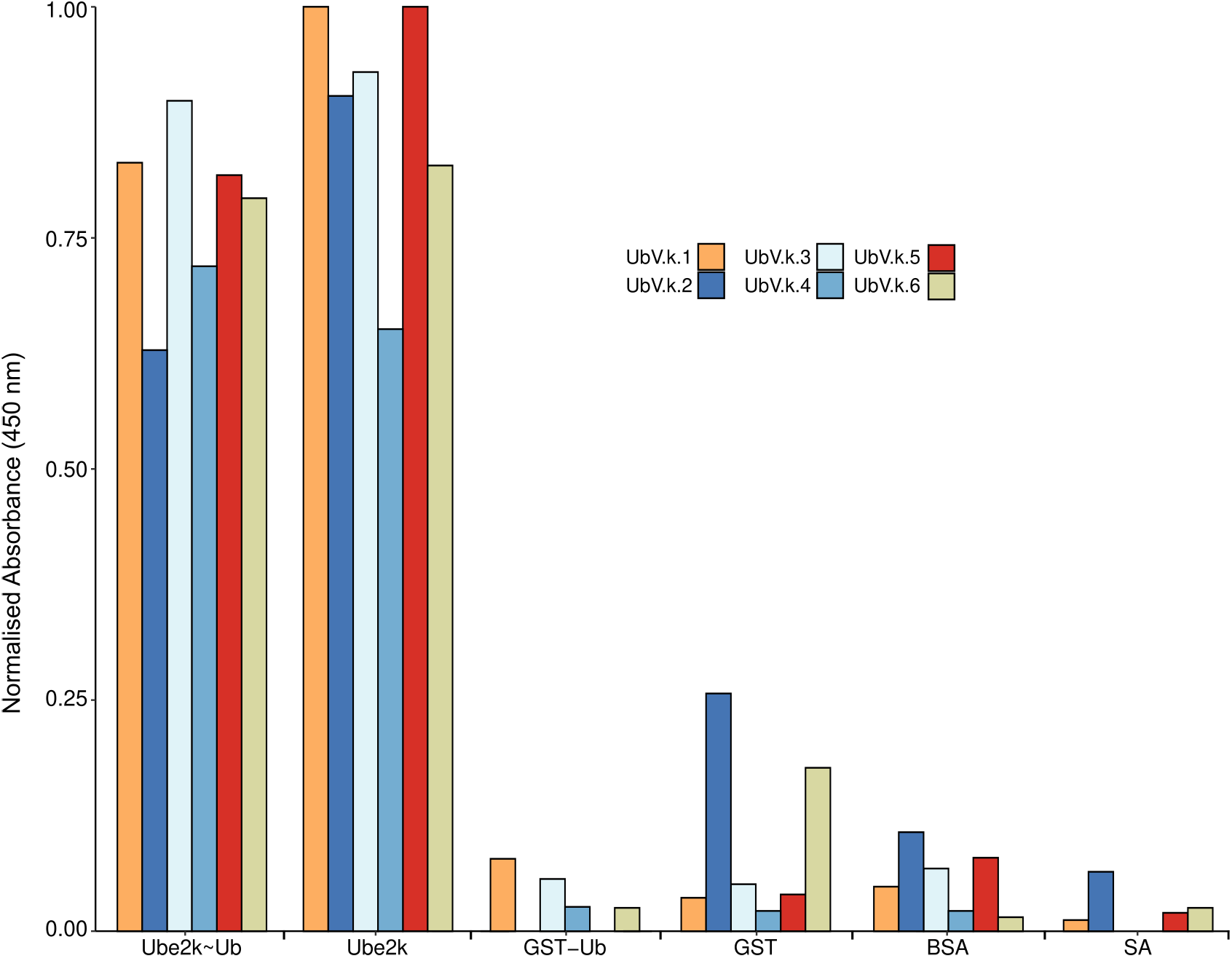
Each UbV binds to Ube2k. The binding of the UbVs to the Ube2k∼Ub conjugate, Ube2k, GST-ubiquitin, and controls was detected using an ELISA as in Figure 2A (n = 1 or 2). GST: Glutathione S transferase; GST-Ub: GST-Ubiquitin; BSA: bovine serum albumin; SA: streptavidin.

**Figure S2.**
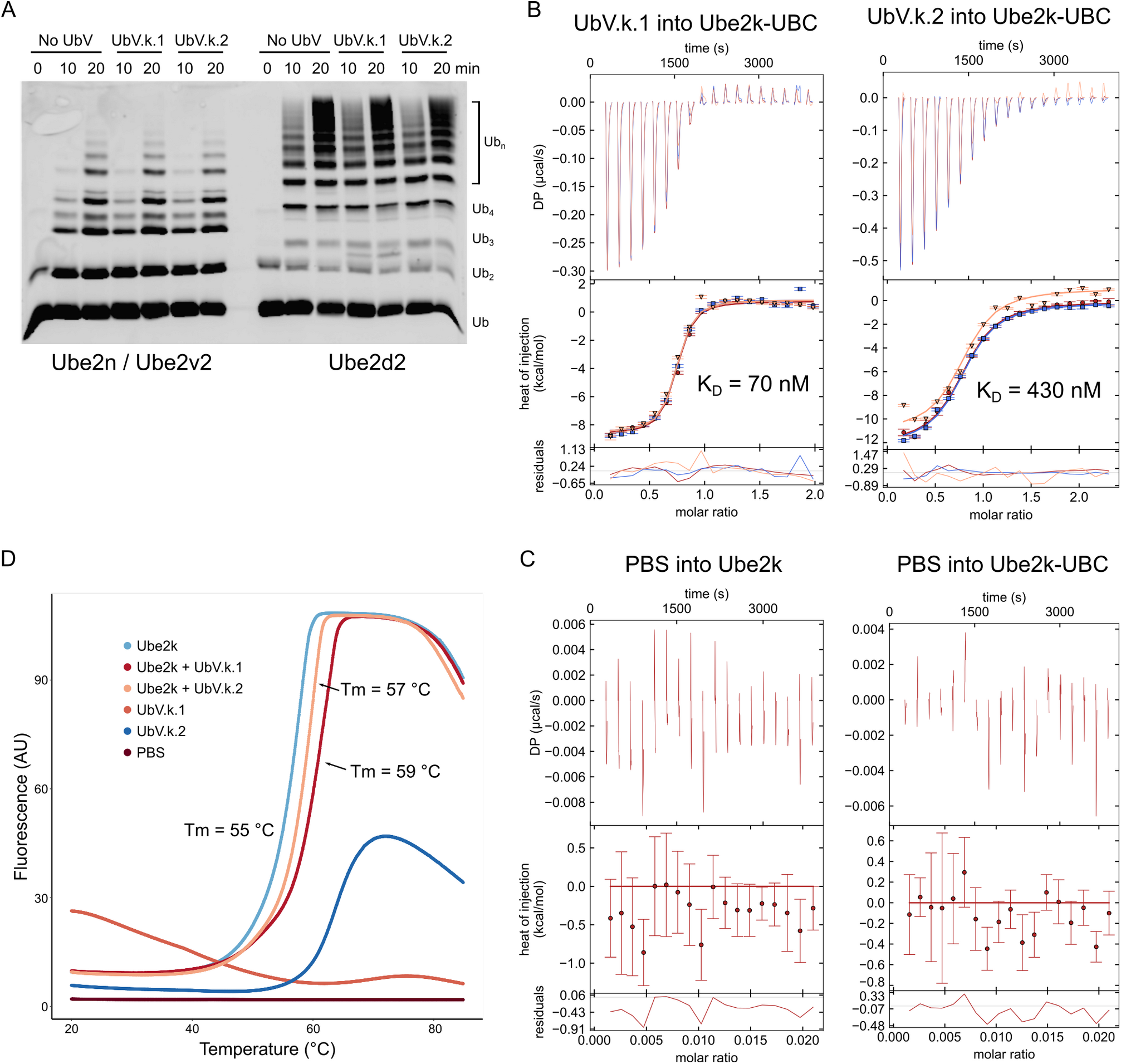
Both UbVs bind tightly and specifically to Ube2k. **(A)** Chain-building assay performed with and without UbV.k.1 and UbV.k.2 with (left) Ube2n/Ube2v2 and (right) Ube2d2. Gel was imaged by fluorescence of Ub-5AIF. **(B)** Thermograms (top) and fitted isotherms (bottom) of ITC measurements performed in technical triplicate (shown in blue, red, and orange) where either UbV.k.1 or UbV.k.2 was injected into Ube2k-UBC. **(C)** Thermograms and isotherms of control reactions where PBS was injected into Ube2k and Ube2k-UBC (n=1). **(D)** Thermal denaturation experiment performed at increasing temperatures with SYPRO orange dye to detect unfolding protein. Performed in technical duplicate. Melting points were calculated with R v4.0.5.

**Figure S3.**
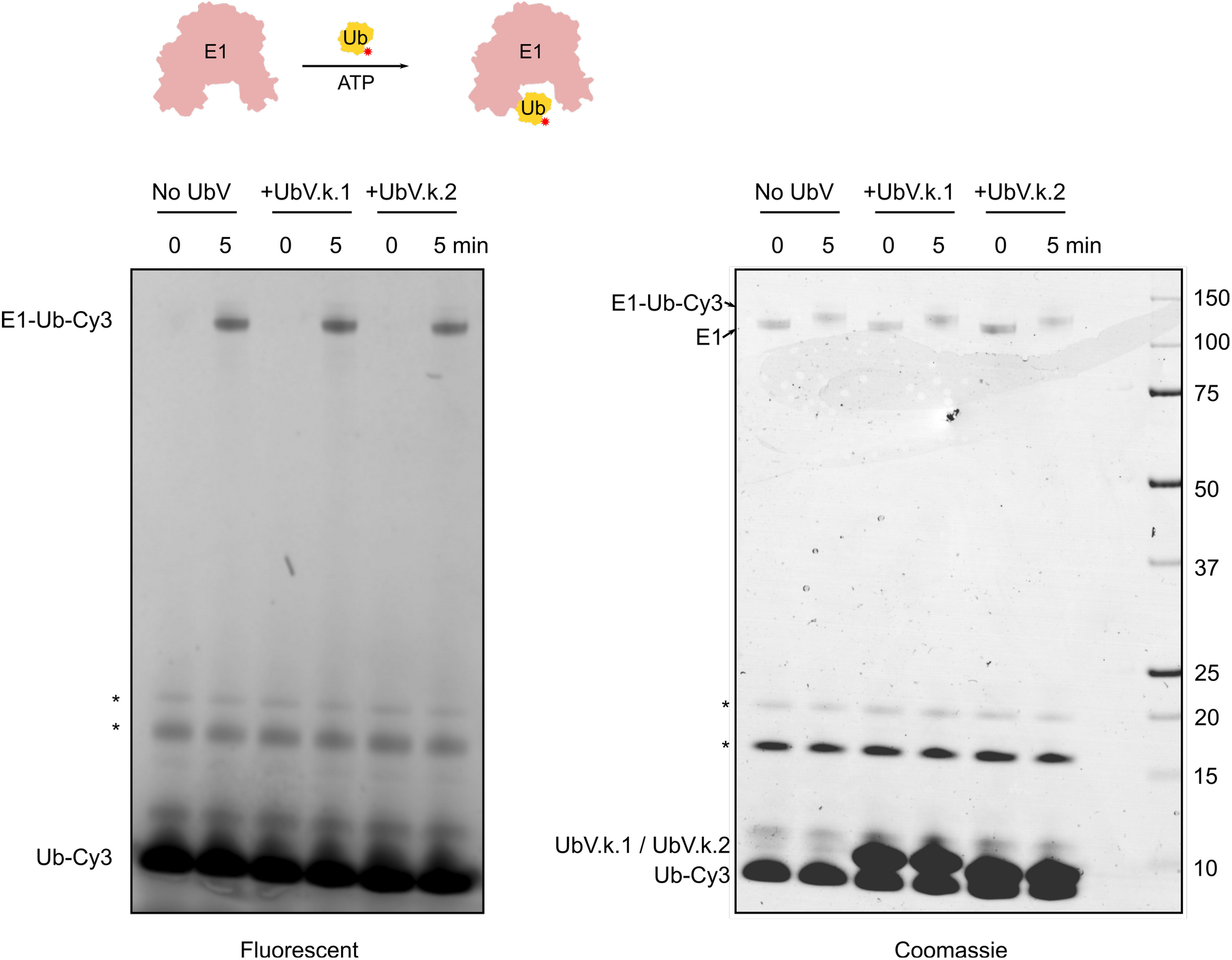
E1 charging assay to assess inhibition by UbVs. (top) Schematic of E1 charging experiment. The red asterisk indicates fluorescent ubiquitin. The ubiquitin activating E1 enzyme was thiolated with fluorescently-tagged ubiquitin with and without UbVs and analysed for fluorescence (left) and Coomassie-stained (right). Asterisks indicate unidentified bands.

**Figure S4.**
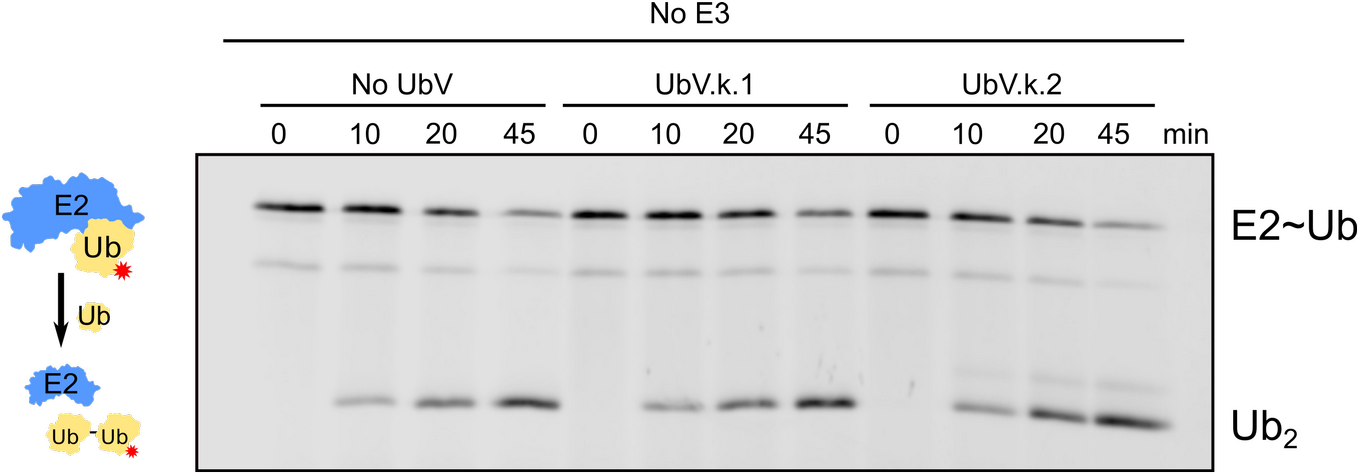
The UbVs do not inhibit basal Ube2k discharge. Fluorescent discharge of a Ube2k∼Ub thioester with and without UbVs. Reaction did not contain an E3 ligase.

**Figure S5.**
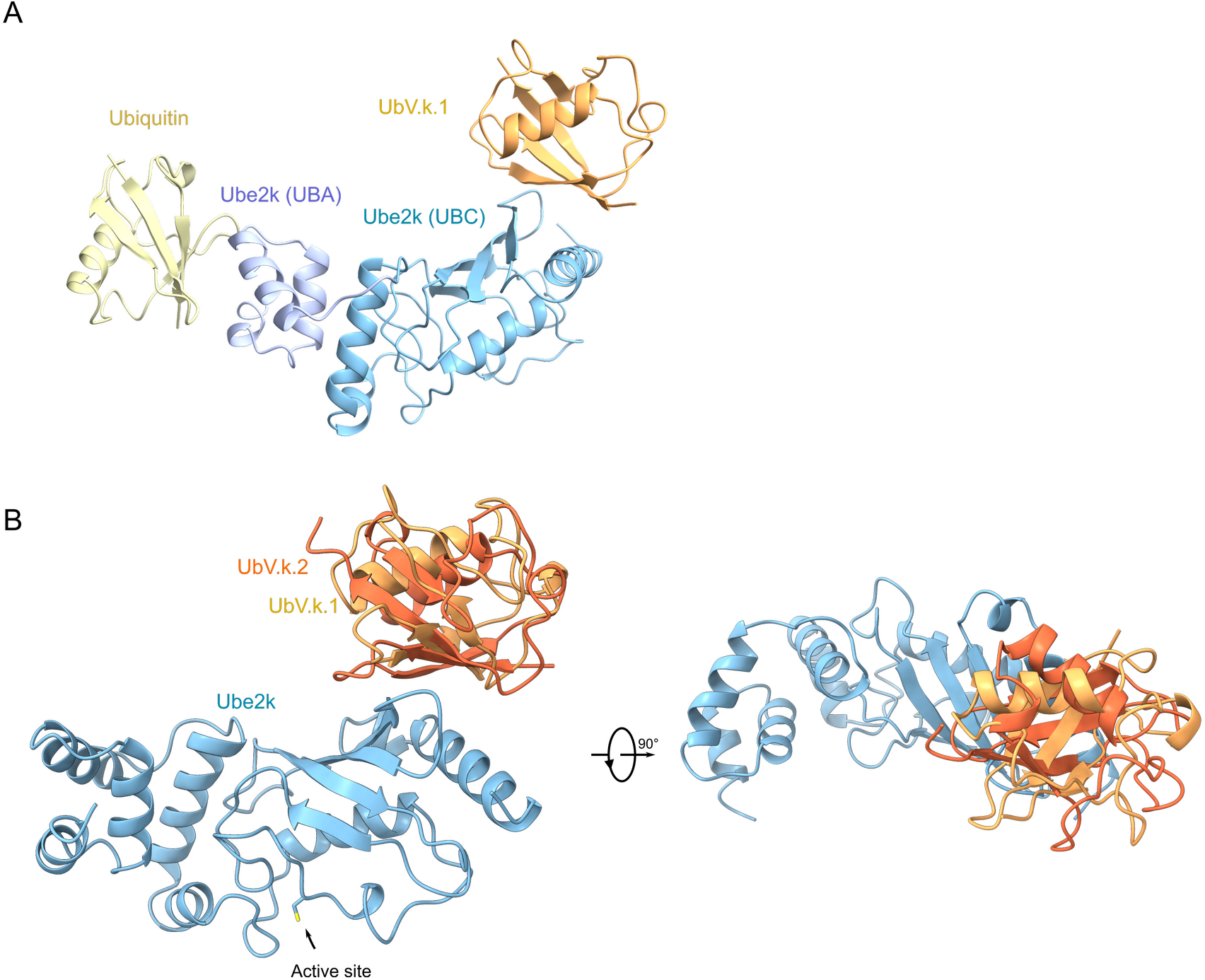
Analysis of the crystal structures of the UbV-Ube2k complexes. **(A)** Overall structure of UbV.k.1-Ube2k∼Ub shown in ribbon representation. The conjugated ubiquitin packs against the UBA domain of a symmetry mate in the crystal. **(B)** Cartoon representation of Ube2k with semi-transparent surface. UbV.k.1 and UbV.k.2 are shown as cartoons interacting with the same surface of Ube2k. The active site is indicated with the Cys shown as a stick. Sulfur atom is coloured yellow.

**Figure S6.**
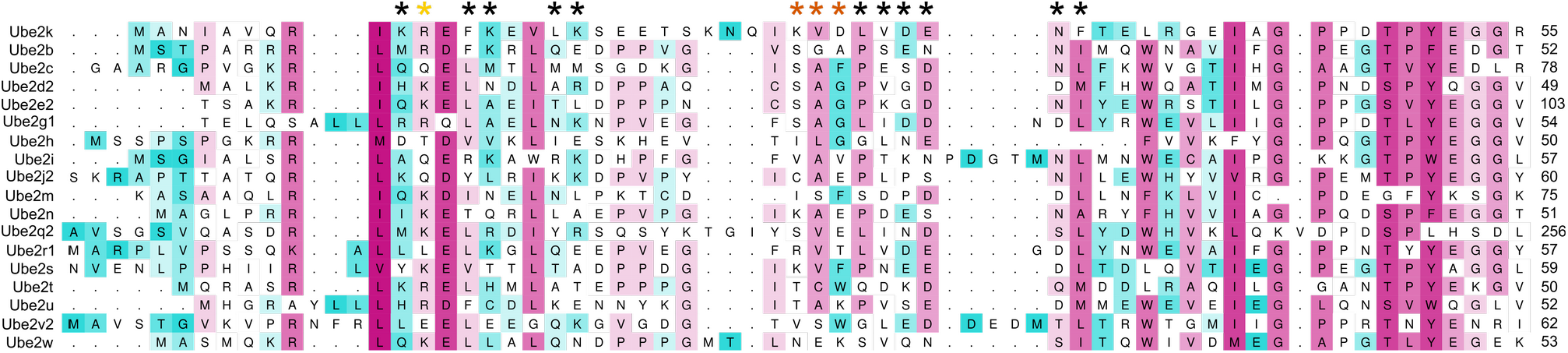
The residues on Ube2k that contact the UbVs are not highly conserved. Sequence alignment of the E2 enzymes used in the ELISA screen (Figure 2A) where maroon indicates high conservation and cyan low conservation. Residues from Ube2k that interact with the UbVs are indicated with asterisks. Black asterisks show overlapping contact residues, while yellow and orange indicate residues contacted only by UbV.k.1 and UbV.k.2, respectively.

## Notes

### Competing Interest Statement

The authors have declared no competing interest.

